# Adaptive optical third-harmonic generation microscopy for in vivo imaging of tissues

**DOI:** 10.1101/2024.05.02.592275

**Authors:** Cristina Rodríguez, Daisong Pan, Ryan G. Natan, Manuel A. Mohr, Max Miao, Xiaoke Chen, Trent R. Northen, John P. Vogel, Na Ji

## Abstract

Third-harmonic generation microscopy is a powerful label-free nonlinear imaging technique, providing essential information about structural characteristics of cells and tissues without requiring external labelling agents. In this work, we integrated a recently developed compact adaptive optics module into a third-harmonic generation microscope, to measure and correct for optical aberrations in complex tissues. Taking advantage of the high sensitivity of the third-harmonic generation process to material interfaces and thin membranes, along with the 1,300-nm excitation wavelength used here, our adaptive optical third-harmonic generation microscope enabled high-resolution in vivo imaging within highly scattering biological model systems. Examples include imaging of myelinated axons and vascular structures within the mouse spinal cord and deep cortical layers of the mouse brain, along with imaging of key anatomical features in the roots of the model plant *Brachypodium distachyon*. In all instances, aberration correction led to significant enhancements in image quality.

## 1. Introduction

Nonlinear microscopy is a powerful tool in the biosciences, enabling non-invasive three-dimensional imaging of living biological tissues, at depths surpassing the capabilities of most linear optical microscopy methods. Nonlinear fluorescence microscopy techniques, such as 2-photon [1,2] and 3-photon [3,4] fluorescence microscopy, often rely on exogenous fluorescent agents for contrast. These methods take advantage of the fact that a fluorescence molecule can be excited to an electronic excited state through the simultaneous absorption of two or more photons, with the subsequent emission of a fluorescence photon occurring when the molecule returns to its ground electronic state. Label-free nonlinear imaging modalities, on the other hand, derive their contrast from intrinsic properties like structural and chemical attributes of the tissue, providing essential information without the need for external labelling agents.

Third-harmonic generation (THG) microscopy is an example of a label-free microscopy approach, relying on a third-order nonlinear optical process [5,6]. This process entails the coherent conversion of three incoming excitation photons into a single photon emitted at exactly three times the energy, resulting in a wavelength of one-third that of the incoming photon. In the case of a focused Gaussian beam, THG signals originate from heterogeneities in the refractive index or third-order nonlinear susceptibility of materials within the focus. In the context of biological imaging, certain tissues and biological structures have been shown to produce robust THG signals, including lipid-rich structures like myelin, calcified bone, and cell membranes [7,8]. When combined with near near-infrared (NIR) excitation wavelengths, THG microscopy enables imaging deep within scattering tissues. However, as the imaging depth increases, the excitation beam accumulates optical aberrations as it propagates through the tissue. These aberrations result in an enlarged excitation focus and reduced focal intensity, ultimately leading to a decrease in the resolution and signal of THG images.

Adaptive optics (AO) is a powerful technology experiencing rapid adoption and implementation across various types of optical microscopes [9–12]. By measuring and correcting the optical distortions introduced by tissues and imaging systems, AO can restore ideal imaging performance deep within tissues. Among the array of AO methods that have been developed for optical microscopy, indirect wavefront sensing methods, which employ a single wavefront shaping device for both aberration measurement and correction, have proven to be well suited for deep-tissue multiphoton imaging of highly scattering samples. One category of methods based on interferometric focus sensing, involves measuring the scattered electric-field point spread function (PSF) at the focal plane of the microscope objective and determining the corrective wavefront via a Fourier transform. Several implementations with varying levels of optical setup complexity have been demonstrated for this strategy [13–15]. Another category of methods comprises modal approaches, which typically employ simpler optical setups [16–20]. Here, aberration modes (e.g., Zernike polynomials) that extend across the entire wavefront are introduced to the excitation beam using a wavefront shaping device, while images are being captured. After acquiring a predefined number of images and assessing the impact of introducing these known aberrations on specific image metrics (e.g., signal or contrast), the final corrective wavefront is derived. In contrast, indirect zonal approaches divide the wavefront into multiple zones and measure the local wavefront distortions in each zone [21–24]. Compared with modal approaches, zonal methods have been demonstrated to be more effective at correcting for the complex aberrations encountered at larger tissue depths [24,25].

We have recently developed a compact adaptive optics module for multiphoton fluorescence microscopes [24]. This module employs an indirect zonal approach based on frequency multiplexing to measure optical aberrations [23]. Using a single high-speed segmented deformable mirror (DM) for both aberration measurement and correction, our module achieved fast aberration measurement with high-power throughput, polarization- and wavelength-independent operation, and easy integration into multiphoton microscopes. We demonstrated high-resolution 2-photon and 3-photon fluorescence imaging in optically challenging environments within the mouse central nervous system in vivo, including imaging of synaptic features in deep cortical and subcortical regions of the mouse brain, and high-resolution imaging of neuronal structures and somatosensory-evoked calcium activity in the mouse spinal cord at great depths [24].

In this work, we integrated our compact adaptive optics module with a THG microscope, which we call from here on the “AO THG microscope”, to enable high-resolution label-free imaging across diverse samples. To validate the performance of our AO THG microscope, we imaged THG-producing non-biological samples, including gold nanoparticles and glass interfaces, under large artificial aberrations and achieved diffraction-limited imaging performance after AO correction. Taking advantage of the high sensitivity of the THG process to interfaces and thin membranes, along with the NIR excitation wavelengths used here (i.e., 1,300 nm), our AO THG microscope allowed us to perform label-free high-resolution in vivo imaging in highly scattering biological model systems at depth. Examples include imaging of myelinated axons and vascular structures in both the mouse spinal cord and deep cortical layers of the mouse brain, as well as imaging of key anatomical features in the highly scattering tissues of the *Brachypodium distachyon* root.

**2. Materials and Methods**

### 2.1 Microscope description

Figure 1 illustrates a schematic of our custom-built THG microscope, as previously described [24]. The laser source comprised a two-stage optical parametric amplifier (Opera-F; Coherent) pumped by a 40-W 1035 nm femtosecond laser (Monaco 1035-40-40; Coherent) at a repetition rate of 1 MHz. Within the extensive tuning range provided by Opera-F (650-920 nm and 1200-2500 nm), we selected an excitation wavelength of 1,300 nm, with an average output power of ~1.5 W (1.5 µJ per pulse). A homebuilt single-prism compressor [26] was employed to minimize the group delay dispersion and achieve a pulse duration of ~54 fs at the focal plane of the objective as measured by an autocorrelator (Carpe, APE GmbH).

**Fig. 1.**
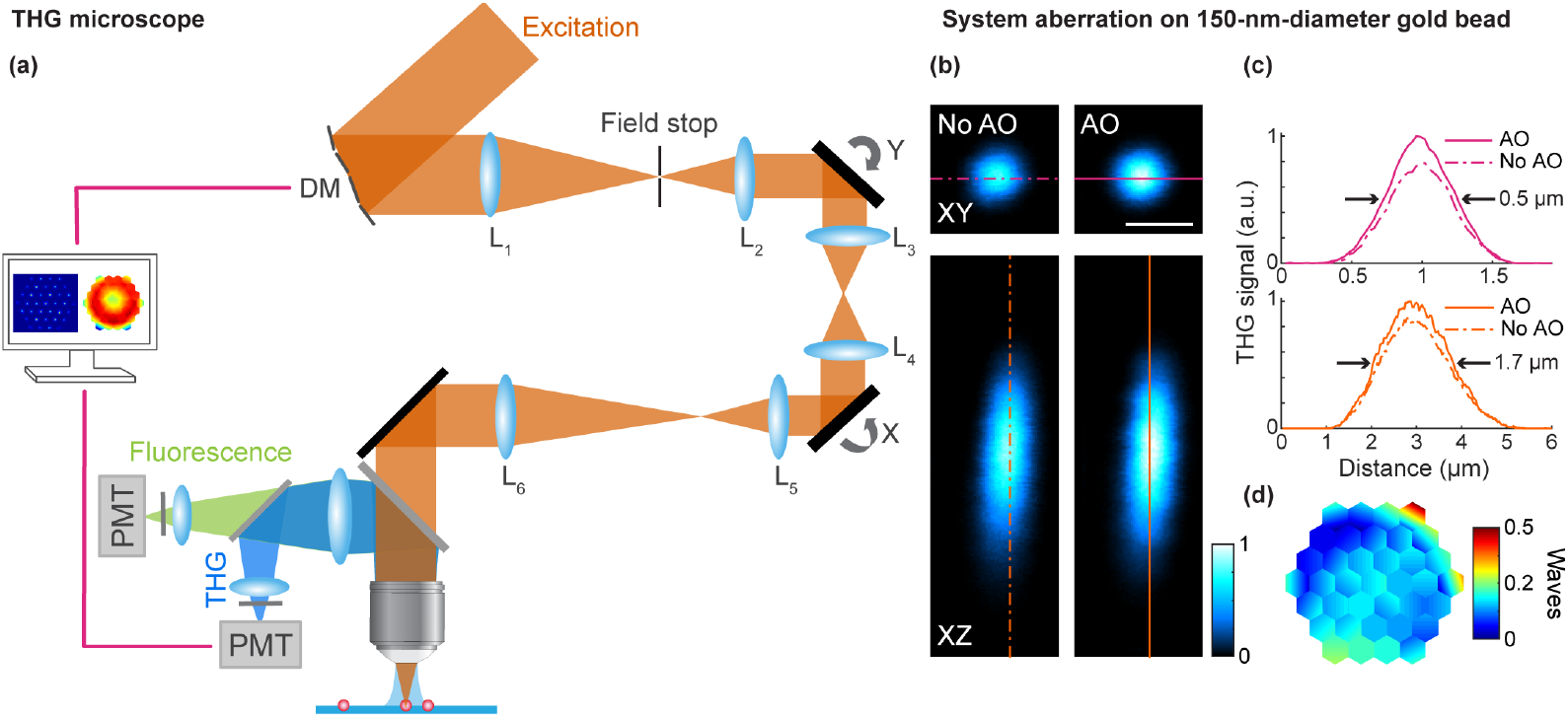
Schematics of the AO THG microscope and system aberration correction. **(a)** Main components of AO THG microscope. DM, deformable mirror; L, lenses; X and Y, galvanometers; PMT, photomultiplier tube. **(b)** Lateral and axial THG images of a 150-nm-diameter gold bead, under 1,300-nm excitation, before and after system aberration correction. Post-objective power: 1 mW. **(c)** Signal profiles along pink and orange lines in (b). **(d)** Corrective wavefront. Scale bar, 1 μm. Microscope objective: NA 1.05 25×.

A deformable mirror (DM) consisting of 37 individual segments (Hex-111-X; Boston Micromachines Corporation) was conjugated to a pair of galvanometers (6215H; Cambridge Technology) used for two-dimensional raster scanning. These galvanometers were optically conjugated to each other and to the back focal plane of a water immersion microscope objective (Olympus XLPLN25XWMP2, NA 1.05, 25×) by three pairs of achromat doublets (Microscope system #1: AC254-100-C and 45-804, AC508-080-C and AC508-080-C, AC508-100-C and SLB-50-600PIR2; Thorlabs, Edmund Optics and OptoSigma. Microscope system #2: AC254-400-C and AC254-300-C, SL50-3P and SL50-3P, SL50-3P and TTL200MP; Thorlabs). The DM image underfilled the back aperture of the objective. For the data presented in Figs. 2a-h, the effective numerical aperture (NA) was ~1.0, corresponding to lateral and axial resolutions of around 0.5 µm and 1.7 µm, respectively (microscope system #1). For all other data presented here, the effective NA was ~0.9, corresponding to lateral and axial resolutions of about 0.6 µm and 2.3 µm, respectively (microscope system #2).

**Fig. 2.**
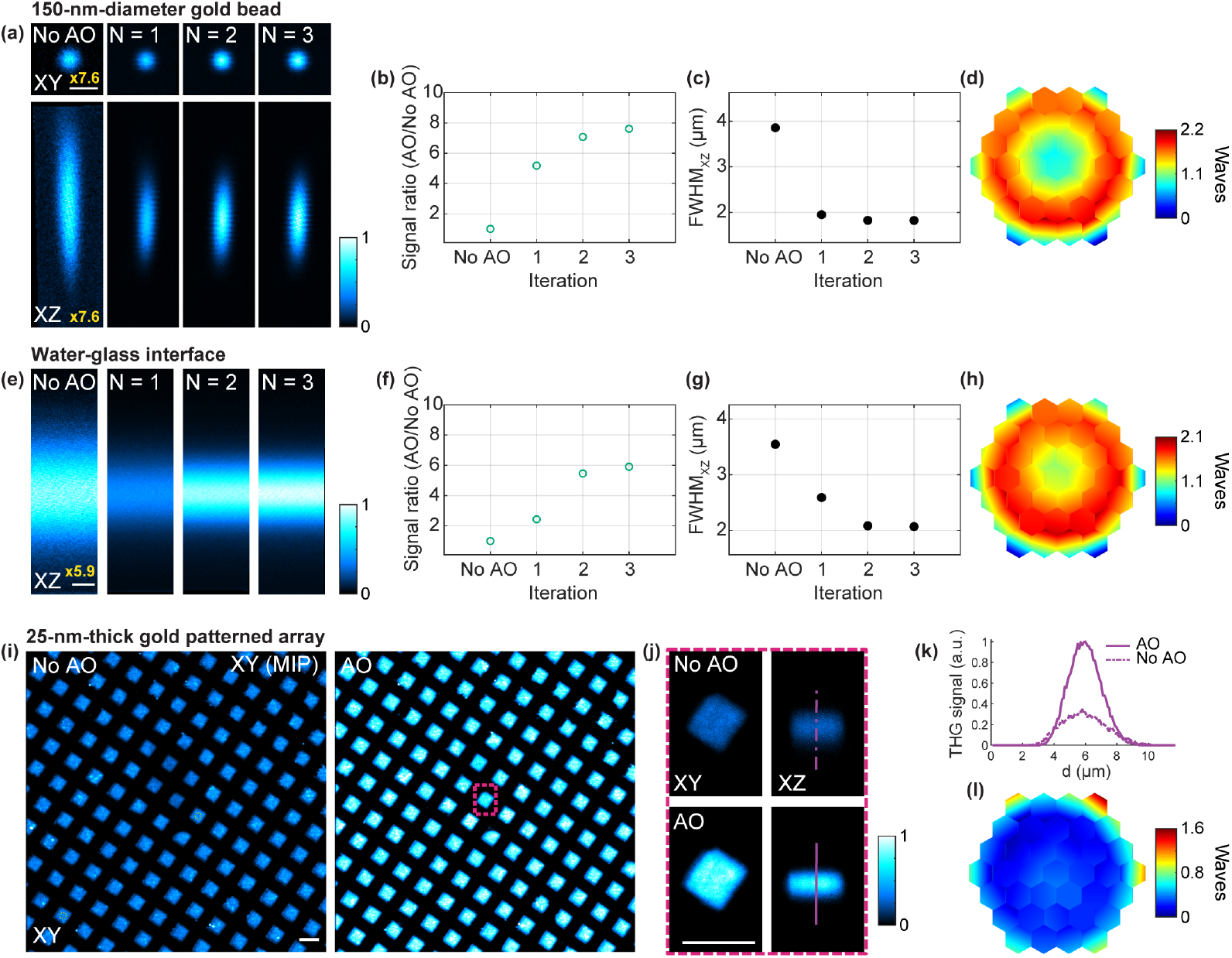
AO improves THG imaging of gold and glass structures under spherical aberrations. **(a)** Lateral and axial THG images from a 150-nm-diameter gold particle with the correction collar of the objective set to 0.17 mm mark, measured without AO correction and after running the aberration measurement a total of N = 1-3 iterations. A 7.6-fold digital gain was applied to No AO images to increase visibility. **(b**,**c)** THG signal improvement (AO/No AO) and axial full width at half maximum (FWHM) of the gold bead as a function of the iteration #, respectively. **(d)** Corrective wavefront after 3 iterations. **(e)** Axial THG images from the top water-glass interface of a #1 coverglass with the objective correction collar set to 0.17 mm mark, measured without AO correction and after running the aberration measurement a total of N = 1-3 iterations. 5.9-fold digital gain was applied to No AO image to increase visibility. **(f**,**g)** THG signal improvement (AO/No AO) and axial FWHM of the air-glass interface as a function of the iteration #, respectively. **(h)** Corrective wavefront after 3 iterations. **(i)** Maximum intensity projections (MIPs) of a 10.5-µm-thick THG image volume of a 2.5-nm-thick gold patterned array sample under a #1.5 coverglass (objective correction collar set to zero), acquired without AO correction and after 3 iterations of aberration measurement. **(j)** Lateral and axial THG images of the region within the red square in (i), acquired without and with AO. **(k)** Signal profiles along the purple lines in (j). **(l)** Corrective wavefront in (i,j). Post-objective power: 0.4, 1.6, and 0.8 mW in (a), (e), and (i,j), respectively. Scale bars, 1 μm in (a) and (e), 10 μm in (i,j). Microscope objective: NA 1.05 25×.

A field stop (iris diaphragm; Thorlabs) was positioned at the intermediate image plane between the DM and the X galvo to block light reflected off mirror segments at large tilt angles. For axial translation of the laser focus, the objective was mounted on a piezoelectric stage (P-725.4CD PIFOC; Physik Instrumente). The THG signal was collected by the same objective, reflected from a dichroic beam splitter (FF665-Di02-25x36; Semrock), spectrally filtered (FF01-433/24-25; Semrock), and detected by a photomultiplier tube (H7422-40 or H10770PA-40; Hamamatsu). Additionally, 3-photon fluorescence signal could be simultaneously detected using a separate channel with a different filter (FF01-680/SP; Semrock). A Pockels cell was used for controlling the excitation power (M360-40; Conoptics).

### 2.2 Aberration measurement method

The details of our frequency-multiplexed aberration measurement procedure were described previously [24]. In summary, we parked the laser focus at one sample location and used the THG signal from this point for aberration measurement. The pupil was divided into 37 regions, corresponding to the number of segments in the deformable mirror (DM). These 37 DM segments were then split into two groups consisting of alternating rows. The aberration measurement procedure first determines the local phase gradients of the wavefront that need to be added to each DM segment so that the beamlets reflecting off them overlap maximally at the focal plane. This was achieved by forming a stationary reference focus using half of the beamlets, while the remaining half of the beamlets were scanned around this focus by applying varying tip/tilt to their corresponding DM segments. At each set of tip/tilt values, we modulated the phase or intensity of the scanned beamlets by varying the piston values of or applying a large tilt to the corresponding mirror segments, respectively, each at a distinct frequency of hundreds of Hz, and recorded the variation in the THG signal. If a scanned beamlet overlapped with the reference focus, modulating its phase/intensity would lead to changes in THG signal, with the amount of overlap determined by Fourier transforming the time-varying THG signal trace and extracting the Fourier magnitude at the corresponding modulation frequency. This enabled the determination of the phase gradients required for maximal overlap and interference between the modulated beamlets and the reference focus, thus providing the tip and tilt angles to be applied to the corresponding DM segment. By swapping the stationary and the scanned DM segments and repeating the phase gradient measurement process, we obtained the local wavefront gradients required to converge all beamlets to the same location in the focal plane. With all the beamlets converging at a common location, the subsequent step of the aberration measurement procedure involves measuring the phase offsets of each beamlet that would allow them to constructively interfere at the focus. This is accomplished by following a similar frequency-multiplexed procedure [27].

Because the starting reference focus is aberrated, both the phase gradient and phase offset measurement procedures may be repeated for a few iterations to achieve optimal aberration correction. With the final corrective wavefront applied to the DM, all beamlets of the excitation light converged and constructively interfered at the focal plane, thus achieving diffraction-limited performance for THG microscopy.

### 2.3 Mouse sample preparation for in vivo brain and spinal cord imaging

All animal experiments were conducted according to the National Institutes of Health guidelines for animal research. Procedures and protocols on mice were approved by the Animal Care and Use Committee at the University of California, Berkeley.

For the in vivo mouse brain imaging experiments, we conducted a chronic cranial window implantation procedure following previously established protocols [28]. Briefly, mice were anesthetized with isoflurane (1–2% by volume in O_2_) and administered buprenorphine (subcutaneous injection, 0.3 mg per kg of body weight). Utilizing aseptic techniques, we created a 3.5-mm diameter craniotomy over the primary visual cortex (V1), leaving the dura intact. A cranial window consisting of a single glass coverslip (No. 1.5, Fisher Scientific) was embedded into the craniotomy and sealed with dental acrylic. A titanium head-post was attached to the skull with cyanoacrylate glue and dental acrylic. Chronic imaging sessions were conducted at least one week following the surgery. During these imaging sessions, mice were immobilized using head fixation and kept under anesthesia with isoflurane (1–2% by volume in O_2_).

For the in vivo spinal cord imaging experiments, we followed an acute window implantation procedure as previously described [29]. Briefly, mice were anesthetized via two consecutive intraperitoneal injections of 1mg/kg body weight urethane, administered 30 minutes apart. Mice were intubated following a tracheotomy to prevent asphyxiation. A dorsal laminectomy was performed at spinal segment T12 after exposing the T11-13 vertebrae, which were stabilized using spinal clamps (STS-A, Narishige). Following a rinse with Ringer solution (135 mM NaCl, 5.4 mM KCl, 5 mM HEPES, 1.8 mM CaCl_2_, pH 7.2), a glass window made of a single coverslip (Fisher Scientific No. 1.5) was placed on the spinal cord along with a custom imaging chamber and stabilized using 2% agarose in Ringer solution. After the surgery, and to ensure adequate hydration during the imaging session, mice were administered subcutaneously with 20 mL/kg body weight of 0.9% physiological saline. Continuous monitoring of blood flow through the central blood vessel was conducted throughout the imaging experiment to ensure tissue health. Animals were euthanized at the end of the imaging session.

### 2.4 Plant root sample preparation

For the in vivo plant root imaging experiments, *Brachypodium distachyon* seedlings were transplanted into microfabricated ecosystem (EcoFABs) containing 50% MS media and allowed to grow hydroponically [30–32]. A glass coverslip was used for the bottom of the EcoFAB so that the roots could be non-destructively imaged. Roots from 3-week-old plants were imaged through a glass coverslip that sealed the bottom of the microfabricated PDMS growth chamber.

## 3. Results and discussion

### 3.1 AO THG microscope characterization and performance evaluation

We first corrected the optical aberrations of the microscope itself (Fig. 1b-d). Applying the corrective wavefront for the optical system aberration (Fig. 1d) led to a ~1.2-fold increase in THG signal of a 150-nm-diameter gold particle (Fig. 1b,c). The lateral and axial full-widths-at-half-maximum (FWHMs) after AO correction were 0.5 µm and 1.7 µm, respectively, consistent with the diffraction-limited resolution of our system. In all following experiments, the images without aberration correction (labeled “No AO”) were acquired after correcting for the aberrations intrinsic to the microscope system.

We first tested our module’s ability to correct for artificial aberrations using the THG signal generated from various samples and found it to enable diffraction-limited imaging performance in all cases (Fig. 2).

For 150-nm-diameter gold particles (A11; Nanopartz) placed under the water-dipping objective, diffraction-limited performance was expected when the correction collar of the objective was set to zero. By rotating the correction collar away from zero to the 0.17 mm mark, we introduced spherical aberrations to the system. For one example gold particle, AO correction of the spherical aberrations increased its THG signal and reduced the axial FWHM of its image (Fig. 2a-d). After the first round of correction, its signal increased by ~5 fold and its axial FWHM decreased from approximately 4 µm to 2 µm. After the second and third rounds of aberration correction, the THG signal continued increasing with an eventual signal gain of ~7.7× (Fig. 2b), and the axial FWHM was reduced to ~1.8 µm. Here and in the examples that follow, only 2-3 rounds of correction were used, as additional iterations did not result in substantial further improvement of the THG signal.

We also corrected for the spherical aberrations introduced by the correction collar using the THG signal of the top water-glass interface of a #1 coverglass (Fig. 2e-h). After 3 rounds of correction, we found a ~6× increase of THG signal, with the axial FWHM of the interface profile reduced from 3.5 µm to 2.1 µm.

Lastly, we applied our AO module to correct for the spherical aberrations introduced by a #1.5 coverglass placed on top of a 2.5-nm-thick gold patterned array sample (here correction collar was set to zero; Fig. 2i-l). After 3 iterations of aberration correction, the THG signal from the gold patterned array increased by ~2.7×. For all three examples here, the corrective wavefronts (Fig. 2d,h,l) indicated spherical aberrations, as expected.

### 3.2 In vivo mouse spinal cord and brain imaging

We next applied our AO microscope to high-resolution in vivo THG imaging of cells and tissue structures in the mouse brain and spinal cord (Fig. 3). A significant source of THG within the nervous system arises from the lipid-rich membranes of myelin sheaths surrounding axons. Consequently, white matter in the brain and spinal cord exhibits strong THG signals. Red blood cells comprise another strong source of THG when using excitation wavelengths near 1275 nm. This phenomenon occurs due to a 3-photon resonant enhancement of the THG signal originating from the Soret transition band in oxy- and deoxy-hemoglobin [33]. In the examples that follow, we used the THG signal from myelin for aberration measurement and found significant enhancements in image quality following AO correction. This enhancement notably improved the visualization of myelinated axons and vascular structures in the mouse cortex and spinal cord.

**Fig. 3.**
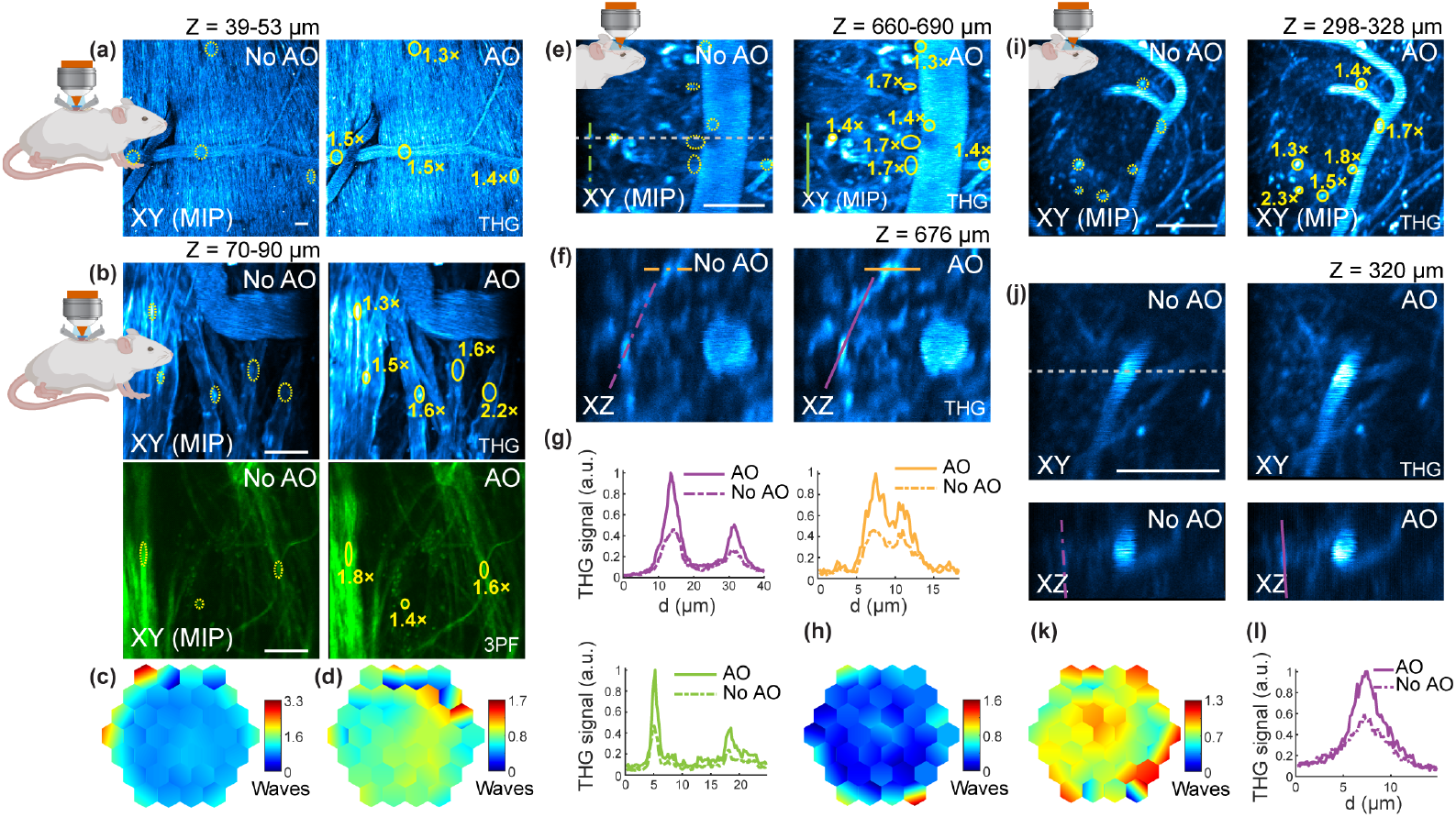
AO improves THG imaging in the mouse spinal cord and cortex in vivo. **(a)** Maximum intensity projections (MIPs) of 14-μm thick THG image stacks (1-μm axial step size) of the superficial layers of the spinal cord in a 31-day-old mouse (C57BL/6J line), under 1,300 nm excitation, without and with AO. Post-objective power: 1.3 mW. **(b)** MIPs of 20-μm thick THG (top) and 3PF (bottom) image stacks of the spinal cord in a 159-day-old mouse (C57BL/6J line, AAV8-Syn-jGCaMP7s), under 1,300 nm excitation, without and with AO. Post-objective power: 2.5 mW. **(c**,**d)** Corrective wavefronts in (a) and (b). respectively. **(e)** THG MIP images of vasculature structures and myelinated axons in the brain of a 323-day-old mouse (Rbp4-Cre line), at 660–690 μm below dura, under 1,300 nm excitation, without and with AO. **(f)** Axial THG images, taken along the gray dotted line in (e), 676 μm below dura, under 1,300 nm excitation, without and with AO. Post-objective power: 19.8 mW. **(g)** Signal profiles along the purple and yellow lines in (f) and green lines in (e). **(h)** Corrective wavefront in (e,f). **(i)** MIP THG images of vasculature structures and myelinated axons in the brain of a 322-day-old mouse (Rbp4-Cre line), at 298–328 μm below dura, under 1,300 nm excitation, without and with AO. **(j)** Lateral and axial (taken along the gray dotted line) images of the same structures as in (i), 320 μm below dura, under 1,300 nm excitation, without and with AO. Post-objective power: 7.3 mW. **(k)** Corrective wavefront in (i,j). **(l)** Signal profiles along the purple lines in (j). THG signal improvement in regions delineated by yellow ovals in (a,b,e,i) is indicated. Scale bars, 20 μm. Microscope objective NA 1.05 25×.

In the mouse spinal cord, we used our AO THG microscope to visualize myelinated axons in the superficial layer of white matter along with red blood cells in vessels (Fig. 3a-d). In one example (Fig. 3a,c), the correction collar of the microscope objective was set to zero and therefore aberration correction compensated for both the spherical aberrations introduced by the glass window as well as the aberrations introduced by spinal cord tissues, leading to THG signal improvements ranging from 1.3-1.5×. In another example (Fig. 3b,d), the correction collar of the microscope objective was set to optimally compensate for the spherical aberrations introduced by the coverglass, and as a result aberration correction compensated only for the aberrations introduced by spinal cord tissues. Here, AO led to THG signal improvements of 1.3-2.2× across the imaging field of view. In addition, we simultaneously acquired 3-photon fluorescence (3PF) images of jGCaMP7s-expressing fine neuronal structures, via a separate detection channel. Here, the correction pattern found using the THG signal from myelin led to 3PF signal improvements of 1.4-1.8×.

We also found a significant improvement in image quality during in vivo THG imaging of myelinated axons and vascular structures in the mouse cortex through a cranial window, at 660–690 μm and 298–328 μm below dura (Fig. 3e-l). Similar to the observations in the spinal cord, we found the blood vessels exhibit strong THG signals throughout the vessel cross sections, resulting from resonance enhancement of THG in hemoglobin. Likewise, we found robust THG signals originating from lipid-rich myelin sheaths enveloping nerve fibers. In these examples, the correction collar of the microscope objective was set to optimally compensate for the spherical aberrations introduced by the glass cranial window. Consequently, the improvement in the THG signal and resolution arose solely from AO correction of brain-induced aberrations. Using the THG signal from myelin for aberration correction, we found signal improvements ranging from 1.3-2.3× after AO correction, as demonstrated by the annotated regions delineated by yellow ovals (Fig. 3e,i) and the line profiles traced across axon tracts (Fig. 3g,l). Unlike its incoherent counterparts, for example, three-photon fluorescence where smaller features are expected to exhibit greater signal enhancement [24], the signal improvement in THG depends not only on sample structure but also on the interplay between the axial coherence length and the axial extent of the excitation focus [34,35], which together led to the varying THG signal improvement observed across the imaging field of view. We also observed an increase in the axial resolution after AO, as evidenced by the shorter axial extent observed in small neuronal structures, such as axons tracts, as shown in the example illustrated in Fig. 3l where the axial FWHM of the line profile decreased from 4.6 to 3.5 μm after AO.

### 3.3 In vivo plant root imaging

In addition to the mammalian tissues, we also tested how well our AO module improved the image quality of label-free THG microscopy in highly scattering root tissues. Here, we used *Brachypodium distachyon* [36,37], a monocot grass that has emerged as an important model system due to its genetic tractability and close relationship with grass crops. Compared to *Arabidopsis thaliana*, another well-known model plant with semitransparent roots [29], the roots of *Brachypodium distachyon* are more optically opaque, thus benefit from the long excitation wavelength of THG.

We performed in vivo AO THG imaging of *B. distachyon* roots through a glass coverslip that formed the root chamber of the EcoFAB devices (Fig. 4a). The correction collar of the microscope objective was set to zero. As a result, aberration correction compensated for the aberration introduced by the coverslip and the root tissue. Imaging was performed in the mature zone of the root, where elongated cells possessed strong THG signals at their cell walls. AO improved cell-wall THG signal by ~2-2.5× (Fig. 4b-f). In lateral images (Fig. 4c, zoomed-in views of the area within the dashed box in Fig. 4b), a few micron-sized gaps between neighboring cell walls were only resolvable after aberration correction (Fig. 4d). After applying corrective wavefront (Fig. 4g), we also observed ~2× increase in cell wall signal in both the epidermis layer and the cortex layer of the root in axial images, enabling better visualization of cells in deep layers (Fig. 4e,f).

**Fig. 4.**
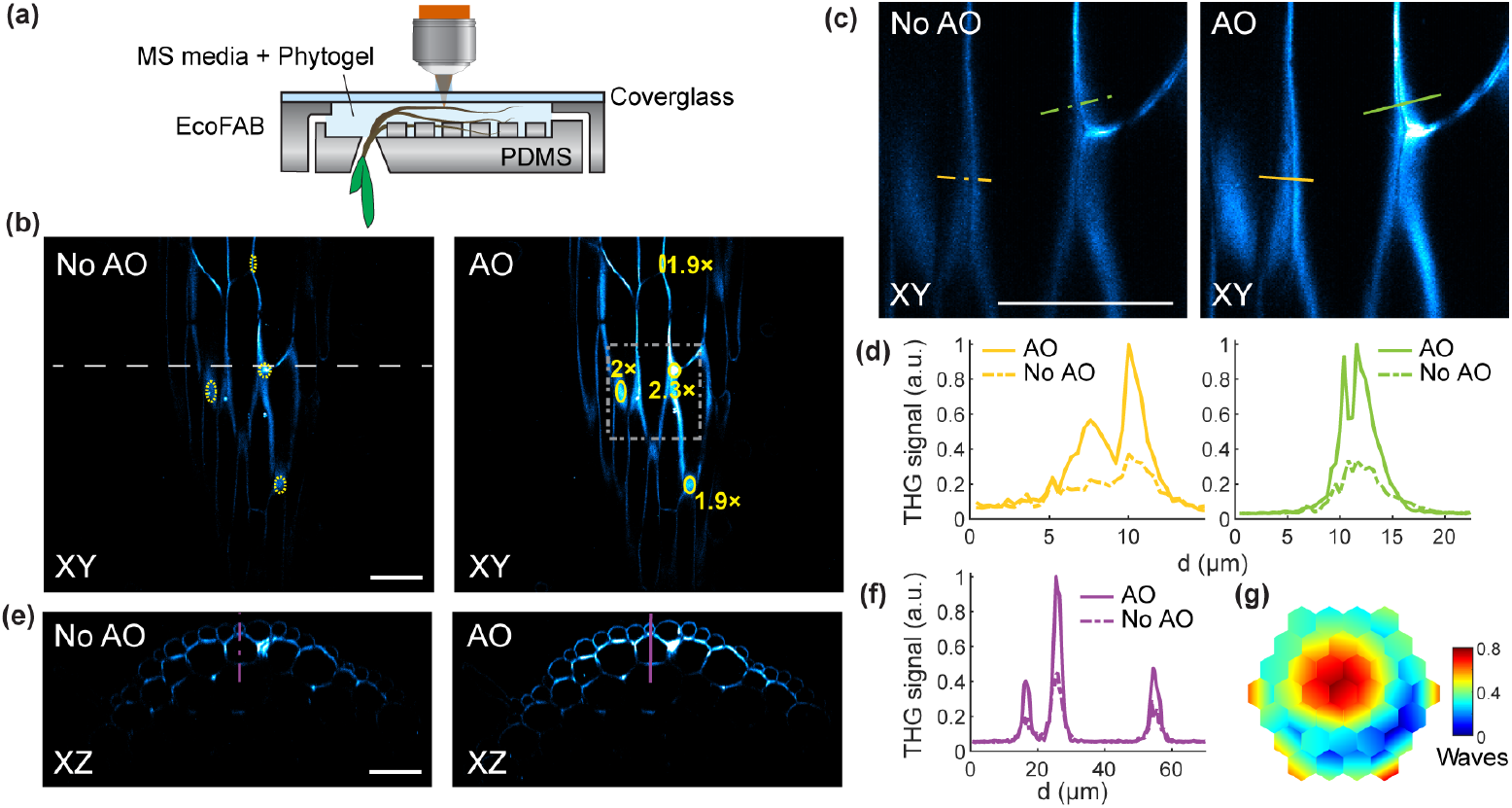
AO improves THG in vivo imaging of *Brachypodium distachyon* root tissues. **(a)** Schematic showing the EcoFAB growth chamber used for imaging the roots of *B. distachyon*. Note that the EcoFAB is inverted for imaging. **(b)** Lateral THG images of the mature zone of a *B. distachyon* root, under 1,300 nm excitation, without and with AO. THG signal improvement along the regions delineated by the yellow ovals is indicated. Post-objective power: 2.2 mW. **(c)** Zoomed-in views of the area in the dashed box in (b). **(d)** Signal profiles along the orange and green lines in (c). **(e)** Axial THG images along the dashed line in (b) acquired without and with AO. Post-objective power: 2.7 mW. **(f)** Signal profiles along the purple lines in (e). **(g)** Corrective wavefront. Scale bars, 50 μm. Microscope objective NA 1.05 25×.

## 4. Conclusion

In this work, we combined a compact AO module with a THG microscope and achieved high-resolution label-free imaging of a variety of biological and non-biological samples. Taking advantage of the exceptional ability of THG microscopy to generate contrast from heterogeneities in specimen optical properties, along with the NIR excitation wavelengths used here (i.e., 1,300 nm), our AO THG microscope allowed us to clearly visualize key anatomical features in highly scattering biological tissues in vivo, without the need for exogenous labelling agents. Examples include the mouse spinal cord, deep cortical layers of the mouse brain, and *Brachypodium distachyon* roots. The image quality improvement achieved after aberration correction was observed consistently across the imaging field of view of hundreds of microns in dimension, consistent with previous studies using fluorescence microscopy [23,24,38,39].

Using bright THG-producing non-biological samples, including gold structures and glass interfaces, our AO module enabled the correction of large amounts of spherical aberration, leading to drastic improvements in the THG signal along with a reduction in the axial extent of the intensity PSF down to the diffraction-limit.

Using our AO THG microscope, we performed label-free subcellular-resolution in vivo imaging of myelinated axons and vascular structures in superficial layers of the mouse spinal cord as well as deep cortical layers of the mouse brain. In both cases, aberration correction led to improvements in the image signal and resolution. In addition, using our AO THG microscope, we performed label-free high-resolution imaging within the highly scattering root tissues of *B. distachyon*. Here, aberration correction led to significant improvements in image quality, greatly improving the visibility of root structures including cell walls. To the best of our knowledge, our results constitute the first demonstration of aberration correction using the endogenous THG signal generated within the previously mentioned biological systems.

We additionally demonstrated that the same correction pattern found using the THG signal resulted in enhancements in the 3-photon fluorescence signal, which was simultaneously detected via a separate channel. In scenarios where the THG signal significantly outweighs any endogenous or exogenous fluorescence signals, conducting the aberration measurement procedure utilizing the THG signal offers advantages. This approach allows for the utilization of lower excitation powers during aberration measurement, a particularly important consideration in highly photosensitive samples such as plant root tissues.

Besides the ability to correct for the optical aberrations introduced by tissues, our AO module effectively compensated for other sources of aberration intrinsic to the experimental setup. In this work, this included the spherical aberration introduced by the glass windows used to gain optical access to tissues: the cranial and spinal windows, as well as the EcoFAB root chamber. Correcting for these sources of spherical aberration becomes especially important when the microscope objective lacks a correction collar. When a correction collar is available, AO can still correct for the additional aberration modes, such as coma and astigmatism, that originate from a tilted window [40].

## Acknowledgement

We thank the Janelia JET team for designing and assembling the dispersion compensation unit and assisting with temporal pulse measurements.

## Data Availability

Data presented in this paper is not publicly available at this time but may be obtained from the authors upon reasonable request.

